# Gamma responses to colored natural stimuli can be predicted from local low-level stimulus features

**DOI:** 10.1101/2023.07.17.549318

**Authors:** Sidrat Tasawoor Kanth, Supratim Ray

## Abstract

The role of gamma rhythm (30-80 Hz) in visual processing is debated; stimuli like gratings and hue patches generate strong gamma, but many natural images do not. Could image gamma responses be predicted by approximating images as gratings or hue patches? Surprisingly, this question remains unanswered, since the joint dependance of gamma on multiple features is poorly understood. We recorded local field potentials and electrocorticogram from two monkeys while presenting natural images and parametric stimuli varying along several feature dimensions. Gamma responses to different grating/hue features were separable, allowing for a multiplicative model based on individual features. By fitting a hue patch to the image around the receptive field, this simple model could predict gamma responses to chromatic images across scales with reasonably high accuracy. Our results provide a simple “baseline” model to predict gamma from local image properties, against which more complex models of natural vision can be tested.

## Introduction

Gamma rhythm (30-80 Hz) in the visual cortex has been related to high level cognition^1, 2^, found to have high stimulus related information^3, 4^ and investigated as a marker for age-related cognitive decline^5^. The dependence of gamma power and center frequency on different features of parametric gratings like contrast, orientation etc. has been well studied^6–11^. Colored stimuli (especially reddish hues) have been discovered to elicit strong gamma responses in the local field potential (LFP) in the primary visual cortex (V1) of non-human primates^12^ as well as in human electrocorticogram (ECoG) recordings^13^. Interestingly, while some studies have found strong gamma responses^14^ to naturalistic stimuli, others have reported weak or inconsistent gamma^15^, which could result from differences in image properties^16^.

This dependence of gamma strength on stimulus features has led to development of image-computable models which predict gamma responses to novel images. For instance, the orientation variance (OV) model^17^ developed using ECoG responses is based on the presence of orientated grating-like features in the receptive field (RF), and the gamma response is proportional to the variance in the outputs obtained by passing the image through predetermined Gabor filters. Other studies have correlated gamma to the presence of large uniform surfaces and the degree of structure^16^ or predictability in an image^18^. Uran and colleagues^19^ predicted RF features from the surrounding image using machine learning, and correlated gamma responses to the degree of predictability. This model used naturalistic stimuli alone whereas the OV model used parametric stimuli. However, it has been argued that an efficient strategy is to build models using artificial stimuli and test them using natural images^20^. We take a three step approach using this strategy to build a simple image-computable gamma model for natural images. The first step is to have a comprehensive knowledge of gamma tuning to as many features (orientation, spatial frequency, color etc.) as possible.

Typically, one feature is studied at a time but when two features are co-varied, the responses offer insights into how they are jointly represented in the brain. The interaction between tuning to different features can be characterized by their separability – to what degree the tuning to one feature is independent of the other. For completely separable interactions, the joint response can be constructed as a function of the individual tunings. Feature independence can also be used to test different theoretical models of tuning^21^. Separability analysis has been carried out for single neurons^22, 23^, membrane potentials^24^ and fMRI responses^25^. Some studies have found spatial frequency (SF) and orientation (Ori) tuning to be separable in monkeys^22^, while others have found interdependence of these features in mice^26^ and cat^27^ visual areas. Surprisingly, separability of LFP responses (specifically, oscillations like the gamma rhythm) has not been well studied although they show preference to Ori and SF^8, 9^. We therefore first investigated the separability of gamma tuning to grating features. Since V1 gamma is also tuned to hues, we extended the separability analysis to HSV (hue, saturation, value) stimuli. Finally, we fit tuning functions to gamma responses to individual features and modelled the overall response as a function of these tuning functions.

The second step involves decomposing a complex natural image into simple features whose response properties can be modelled. From the built models, we could estimate gamma responses to two classes of stimuli. The first are achromatic image patches that can be approximated by spatial Gabors (sinusoidal grating with dark and light bars weighted by a 2- dimensional spatial Gaussian). The second category are color images where the luminance, hue and saturation remain uniform over an area. These can be approximated as uniform chromatic patches in the HSV space.

The final step involves applying the model learnt in the first step to the image parameters approximated in the second to get a prediction of the gamma response to the image.

## Results

Data were recorded using a hybrid array (with 81 microelectrodes and 9 ECoG electrodes) from two awake monkeys (M1 and M2) who performed a fixation task while parametric^28^ or natural images^4^ were presented. Details about the protocols and electrodes can be found in Methods and Table 1.

**Table 1:** Separability index and marginal correlations. Mean separability index (si) for all protocols, averaged over all electrodes for each monkey. The number of electrodes for which the si is significantly different from a random distribution is mentioned in column 3.

| **Protocol type.**<br>**Number of available**<br>**electrodes** | **Mean Separability**<br>**Index,**<br>**$s_i \pm SD$** | **Number of**<br>**electrodes with**<br>**significantly**<br>**different $s_i$.**<br>**t-test $p \leq 0.05$** |
| --- | --- | --- |
| **SF-Ori**<br>**n1= 82**<br>**n2= 21** | M1: $0.98 \pm 0.01$<br>M2: $0.98 \pm 0.01$ | M1: 80/82<br>M2: 21/21 |
| **Size-Ori**<br>**n1= 29**<br>**n2= 22** | M1: $0.97 \pm 0.02$<br>M2: $0.97 \pm 0.01$ | M1: 29/29<br>M2: 18/22 |
| **Con-Ori**<br>**n1= 82**<br>**n2= 21** | M1: $0.95 \pm 0.02$<br>M2: $0.99 \pm 0.01$ | M1: 81/82<br>M2: 21/21 |
| **Hue-Sat**<br>**n1= 65**<br>**n2= 39** | M1R: $0.94 \pm 0.02$<br>M2R: $0.95 \pm 0.01$ | M1R: 52/65<br>M2R: 39/39 |
| **Hue-Val**<br>**n1= 65**<br>**n2= 39** | M1R: $0.97 \pm 0.02$<br>M2R: $0.99 \pm 0.00$ | M1R: 62/65<br>M2R: 39/39 |
| **Sat-Val**<br>**n1 = 19** | M1R: $0.98 \pm 0.01$ | M1R: 19/19 |

### Separability of Gamma Responses

We first tested a pair of stimulus features for separable gamma tuning. If separable, joint responses can be reconstructed from individual feature dependent functions. We followed two methods, singular value decomposition (SVD^24^) and marginals^22^, to analyze the joint tuning as a product of independent factors. SVD factorizes a matrix into independent vectors, whose weighted sum of products can reconstruct the matrix (Equation 1). For perfect separability, the matrix can be reconstructed as a product of the first left and right singular vectors, weighted by the first singular value. We quantified this measure of independence by a separability index (*s_i_*) which is 1 for a perfectly separable case (see Ref^22^ and Equation 2). SVD provides independent factors representing some combination of the 1-dimensional tuning which may not always be the same as the marginals, which are obtained by simply averaging across rows or columns to get the dependence on one feature irrespective of the second feature. Marginals are an explicit and easy way to obtain tuning of the independent features.

For achromatic gratings, we considered Ori, SF, contrast (Con) and size as the features. Since the number of stimuli required to investigate joint tuning along all features was prohibitively large, we fixed one feature (Ori) and investigated its joint tuning with other features (SF, Con and Size).

#### SF-Ori

Full-screen, maximum contrast gratings varying in Ori (8 values) and SF (5 values) were displayed in the “SF-Ori” protocol (see Methods). Fig. 1A shows the change in power from baseline to SF-Ori stimuli for a typical electrode from M1. These were obtained by estimating the power between 250-500ms after stimulus onset and normalizing by the baseline power (-250-0ms; see Methods). These stimuli evoked strong gamma rhythms as can be observed by the spectral peaks especially for SF of 2 and 4 cycles per degree (cpd) stimuli, particularly at 90° orientation (cyan traces). These large stimuli elicited two peaks, which have been termed slow and fast gamma^9^. As has been previously reported using the same monkeys^28^, gamma peak frequency varied across SF-Ori conditions (as well as size and contrast manipulations, see Ref^28^). As shown later, natural images often did not induce a strong gamma rhythm and for most, a clear peak was not present in the spectrum. Therefore, we considered a fixed frequency range of 30-80Hz (grey vertical lines in Fig. 1A) and used the change in power averaged in this range as our “gamma response”. This is shown as a 2-dimensional matrix over SFs and Oris in Fig. 1B (scaled to have an area of 1 to be represented as a joint density function), over which we performed SVD and marginal analysis.

**Figure 1:**
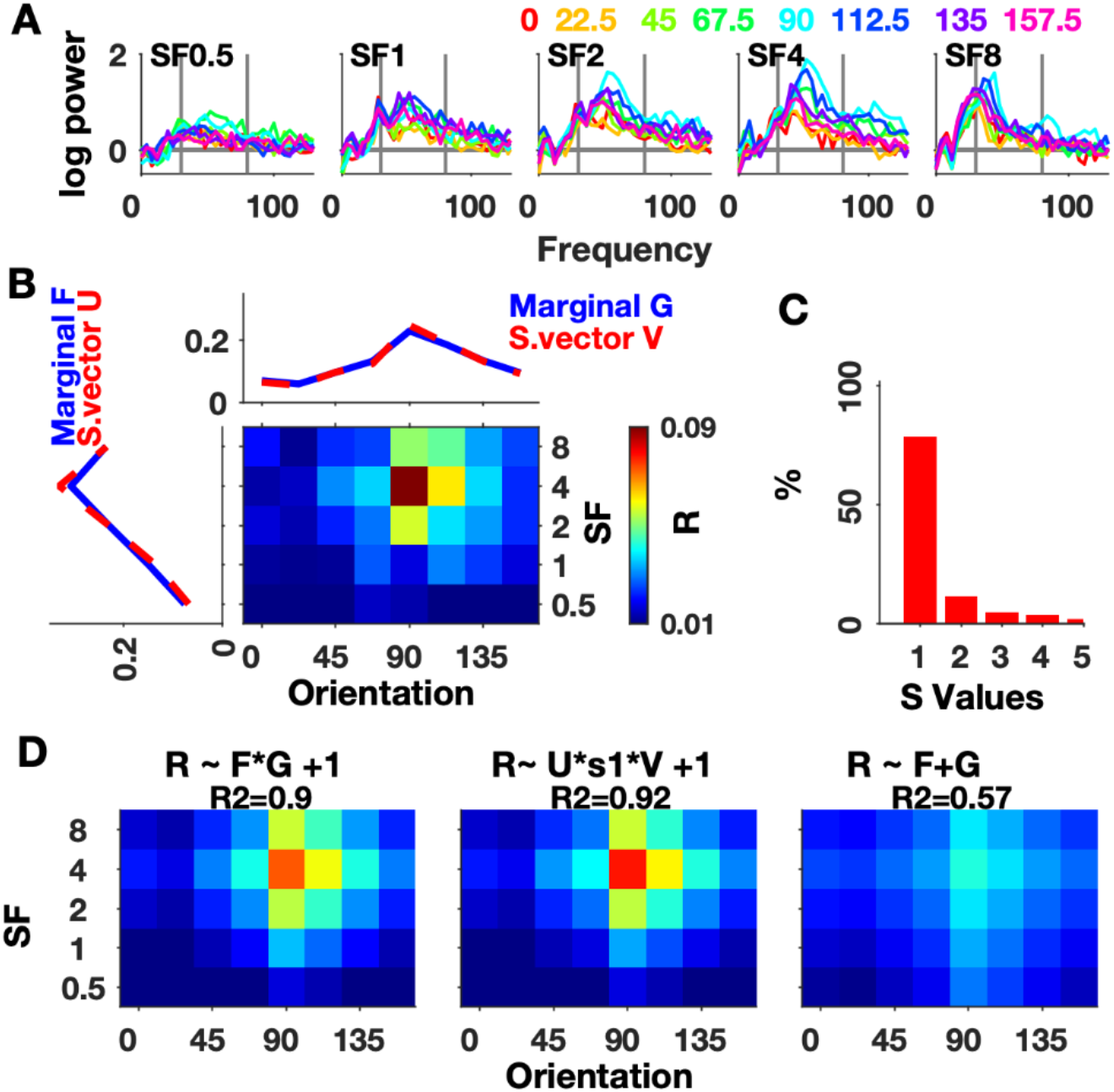
Separability of orientation and spatial frequency tuning for one electrode (M1, e7). (A) Frequency spectra of the log change in power from baseline (-250 – 0 ms) in 250-500 ms period for 5 spatial frequencies (SF) and 8 orientations (Ori) for the SF-Ori protocol. Colors correspond to orientations as indicated on top of the figure. Gray vertical lines mark the chosen gamma band (30-80 Hz). (B) Trial averaged gamma responses (30-80 Hz) to stimuli varying in Ori (columns) and SF (rows) for this electrode. The row and column marginals are shown in blue. The red curves show the first singular vectors from the SVD decomposition of the 2D matrix. Responses, marginals and singular vectors are scaled so that area under the curve equals 1. (C) The percentage contribution of ordered singular values obtained from SVD. (D) Modelled separable joint matrix as a function of independent vectors with the corresponding ratio of variance explained (R^2^) by product of marginals (left), by product of singular vectors (middle) and by sum of marginals (right).

The first singular value represents the contribution of the first singular vectors from the SVD (red curves in Fig. 1B) to the joint distribution. The relative contribution of the singular values (Fig. 1C) reveals that the first one had a much higher contribution compared to the others, and consequently the joint distribution had a high separability index (*s_i_*=0.97). Based on the product of the first singular vectors, we modelled a joint response matrix (Fig. 1D middle, see Methods). We obtained the coefficient of determination (R^2^=0.92) which quantifies how much of the data variance can be explained by the model. In the second approach, we modelled the joint response as a product of the marginal Ori and SF tunings which are plotted in Fig. 1B (blue curves) and were highly overlapping with the first singular vectors. The modelled 2-dimensional response using the product of marginals (Fig. 1D, left) also had a high R^2^ (0.9). For comparison, we also used an additive model (see Methods) which is another separable marginal model. It explained much less variance (Fig. 1D, right; R^2^=0.57) compared to the product models.

This behavior was mirrored across microelectrodes and ECoGs. The average separability index was 0.98±0.01SD (n=82) for M1 and 0.98±0.01SD (n=21) for M2. We tested *si* against bootstrapped values (see Methods) and found it to be significantly different (t-test, p≤0.05) from separability indices of randomly constructed matrices for 80 (of 82) electrodes of M1 and all electrodes of M2. Average R^2^ for the SVD based model (Fig. 2, “SF-Ori” plot; green bars) was 0.83±0.13SD (M1) and 0.87±0.08SD (M2), similar to values obtained by the marginals product model (magenta bars): 0.82±0.13SD (M1) and 0.87±0.08SD (M2). The additive model (blue bars) explained significantly lesser variance: 0.64±0.11SD (M1) and 0.78±0.11SD (M2).

**Figure 2:**
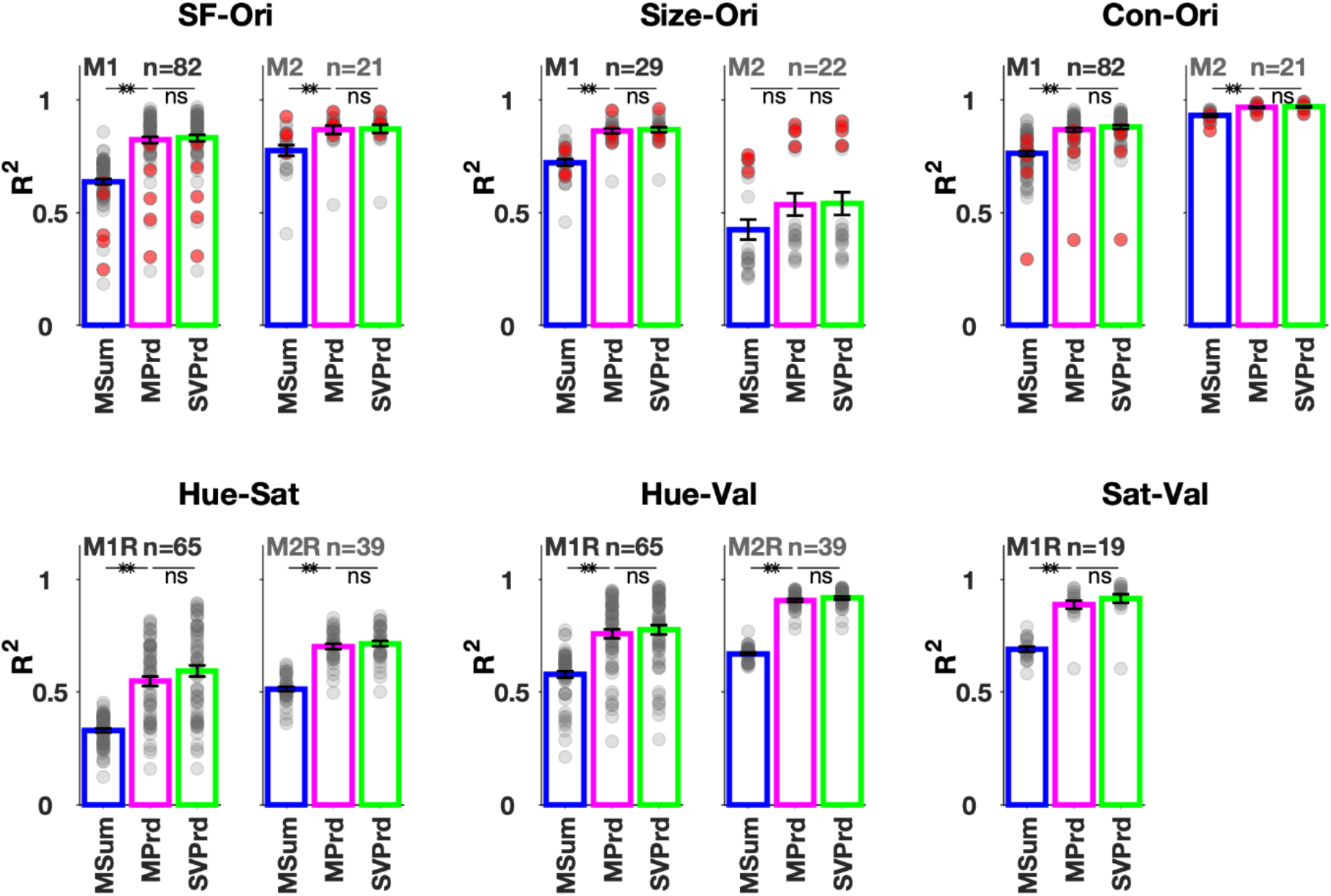
Population Level Separability. The percentage of variance explained (R^2^) of all protocols. M1R and M2R represent the right hemispheres, while M1 and M2 are the left hemispheres of the same two monkeys. The grey and red dots represent the individual microelectrodes and ECoGs respectively. The electrode averages are plotted as bars for the three approaches used – sum of marginals (MSum, blue), product of marginals (MPrd, magenta) and product of singular vectors (SVPrd, green). (**: p<=0.005, unpaired t-test).

#### Size-Ori and Con-Ori

Responses to varying size and orientation (Size-Ori) and varying contrast and orientation (Con-Ori) were recorded separately. In the Size-Ori protocol, full contrast gratings at 8 orientations and 6 sizes were displayed centered on specific RFs (see Methods). The Con- Ori protocol used full-screen gratings at 6 (or 7) contrasts and 8 (or 4) orientations (see Methods). As before, we obtained the 2-dimensional gamma responses (power change in 30- 80Hz) for these protocols for all available electrodes, derived the marginals and singular vectors from them, and made marginal product, marginal sum and singular vector product based models. Their R^2^ values and other metrics are summarized in Fig. 2 and Table 1.

SVD analysis revealed high separability of the 2-dimensional responses with separability indices close to 1 (Table 1) for both Size-Ori and Con-Ori protocols. The product models explained significantly more variance than the additive model (Fig. 2), but the R^2^ for the two product models were not significantly different from each other (statistics in Fig. 2). While the separability index was high for all electrodes, the explained variance was occasionally not very high, especially for Size-Ori protocol for M2. This was simply because the microelectrode RFs were very foveal for M2, and therefore the response saturated quickly and did not increase monotonically with size (see Fig. 3A for gamma versus size; see Fig. 3G- H of Ref^28^ for change in power spectra). Indeed, ECoGs (red points) had higher R^2^ values since their RFs were not foveal. However, even for this protocol, the multiplicative model performed much better than the additive one.

**Figure 3:**
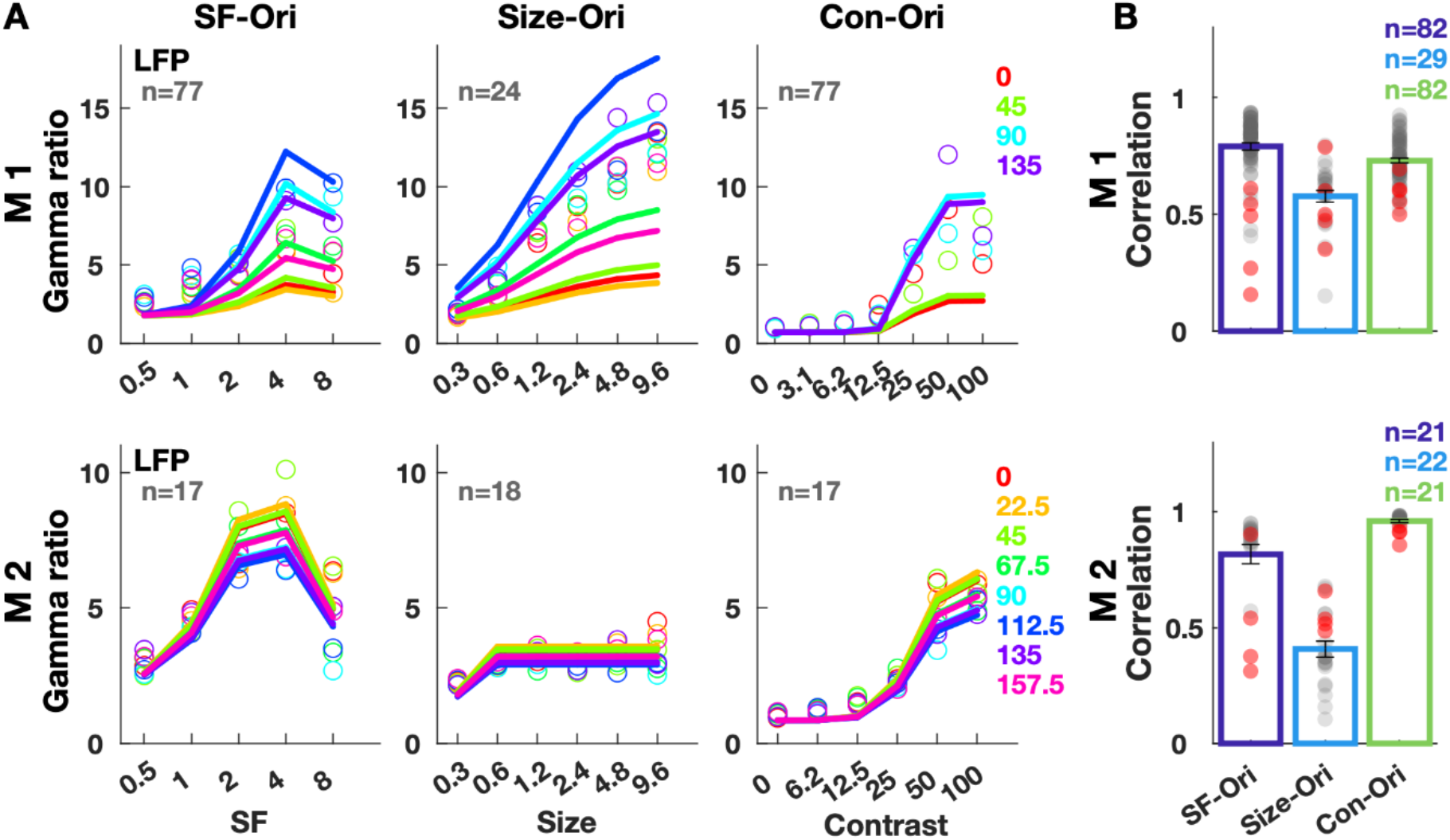
Actual gamma response and modelled responses. Response refers to the change in gamma band (30- 80) power from baseline (-250-0ms) in 250-500ms period. (A) Data (open circles) and estimated fits of responses to the SF-Ori, Size-Ori and Con-Ori stimuli, averaged across electrodes and 2 folds of a single cross-validated iteration. Colors correspond to orientations. (B) Average correlation across electrodes between actual and estimated responses (bars). Grey circles denote correlation values for individual microelectrodes and ECoGs are shown in red. The correlation values of each electrode are averaged over 3 iterations of cross-validation. Error bars denote SEM.

#### Hue, Saturation and Value

Responses to parametric chromatic stimuli were recorded from the right hemispheres of the same monkeys in a different set of experiments^12^, and are labelled as M1R and M2R. We tested separability using full screen protocols in which two features out of Hue, Saturation (Sat) and Value (Val) were varied. Hue-Sat protocol displayed 6 unique hues at 5 Sat levels, while the Hue-Val protocol used 6 hues with 5 luminance levels. Sat-Val protocol (only for M1R) used 5 levels of both saturation and value for red hue (see Methods). For all protocols, we constructed a 2-dimensional matrix for each electrode from the mean change in power in the gamma range, and then performed separability analysis. The results are summarized in Table 1 and Fig. 2. The marginal additive model explained the least variance as with grating stimuli. R^2^ values in Fig. 2 indicate that the first singular vectors could explain most of the data variance which was not significantly different from the marginal product model (statistics in Fig. 2). In the Hue-Sat protocol of M1R, average R^2^ was lower than the other protocols, likely because at low saturation, responses were not well separable across hues for some electrodes (Fig. 4A, “Hue-Sat” plot, open circles) since the stimuli simply were white/dimmed irrespective of hue. Nonetheless, the main results – superior performance of product models compared to additive model, and comparable performance of marginal and SVD based models – were observed for this case also.

**Figure 4:**
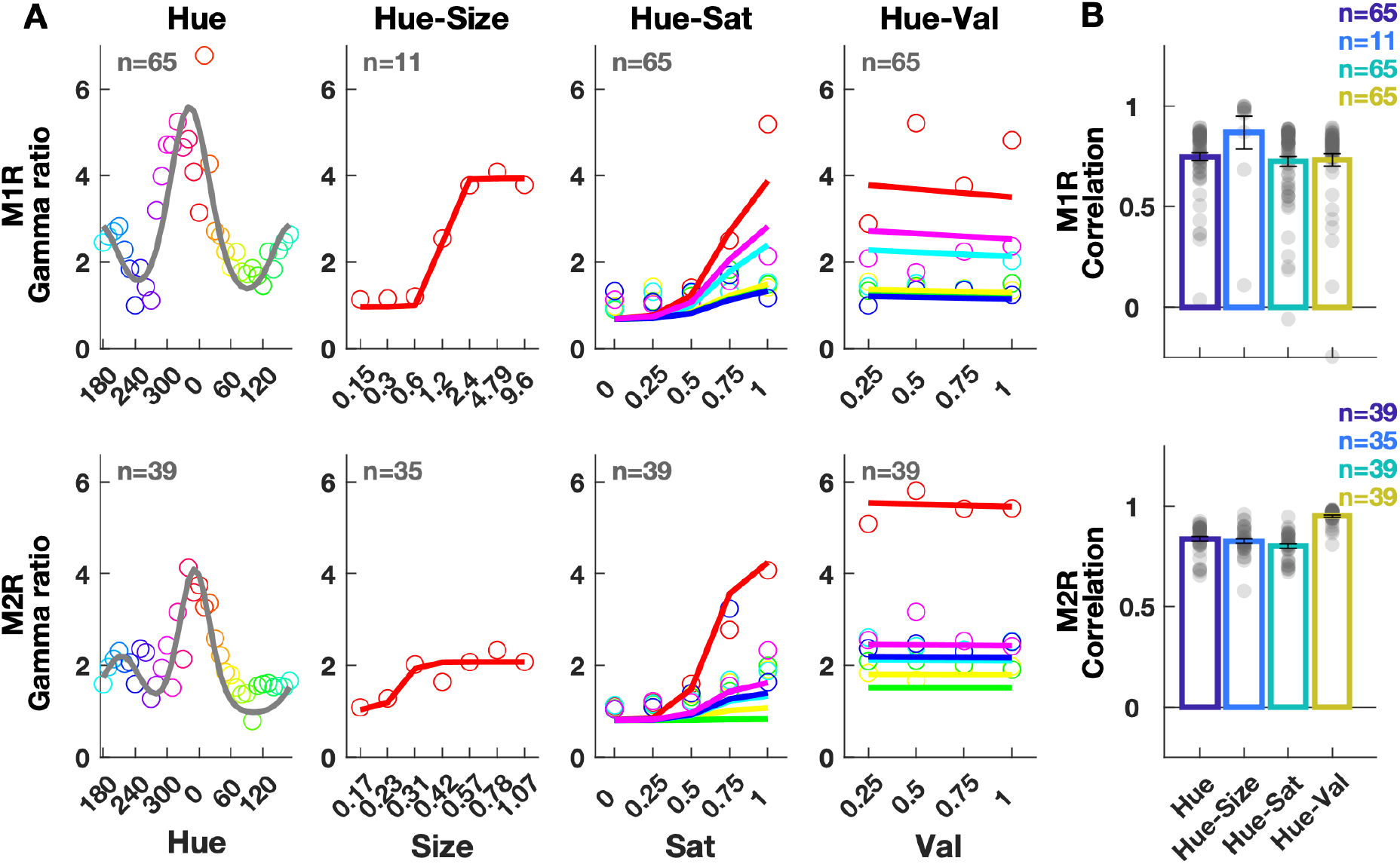
Gamma responses to hue stimuli and estimates. Response refers to the change in gamma band (30- 80) power from baseline in 250-500ms period. (A) Actual response (open circles) and estimated fits of responses to the Hue, Hue-Size, Hue-Sat and Hue-Val stimuli, averaged across 2 folds of a single cross-validated iteration for all microelectrodes. (B) Correlation between actual and estimated responses averaged across electrodes (bars). Grey circles denote correlation values for individual electrodes. The correlation values of each electrode are averaged over 3 iterations of cross-validation. Error bars denote SEM.

Overall, the separability results indicated that 2-dimensional responses to parametric stimuli were largely separable and expressed as products of independent functions. These functions could be estimated from the 1-dimensional tuning to individual features, as discussed next.

### Modelling of responses

We modelled the independent feature tuning of gamma by functions reflecting the shape of the gamma tuning curves, using the same data as for the separability analysis. We did not systematically compare different function classes but chose simple functions popularly used for modeling such responses. We then extended the pairwise separability to all features and expressed the response as a multidimensional product.

#### Grating responses

We modelled the SF tuning as a Gaussian function on the SF values, with a center and a spread parameter (Equation 6), and the orientation tuning as a von Mises function over the angular space (Equation 7). Size and contrast were considered as scaling functions which were maximum at the full-screen size and 100% contrast respectively and were modelled as sigmoidal functions (Equations 8 and 9). Each of these functions was described by two free parameters (see Methods).

Since the features – SF, contrast and size – were separable from orientation, we extended this finding and expressed the response as a product of 4 individual functions (Equation 10A; 8 free parameters characterizing the centers and spreads for Gaussian and von Mises functions and the slopes and midpoints for the sigmoidal size and contrast functions).

We first performed a parameter estimation procedure for each electrode separately, where trials from SF-Ori, Size-Ori and Con-Ori protocols were divided into two folds, one of which was averaged to yield the gamma responses to calculate the model parameters. Specifically, to get the SF parameters, SF-Ori responses were pooled across orientations and fit to equation 6 using least squares fitting. Similarly, responses were pooled across SFs to fit orientation parameters (Equation 7), and so on. Subsequently, the predicted gamma response from the model was compared against the actual response obtained by trial-averaging the second fold.

Supplementary Figure 1 shows the electrode-wise parameters. As also reported in previous studies^9, 28–30^, the parameters tended to cluster together allowing for a major simplification: we used a common set of parameters for all electrodes of each monkey by choosing the medians of these distributions. Therefore, all electrodes used the same shape of tuning functions resulting in a common estimated value (Equation 10A). Electrode estimates could therefore only vary by gain and offset terms (Equation 10B).

Fig. 3A displays the actual gamma response (open circles) and the estimates (lines; using Equation 10B) averaged across microelectrodes. Despite using the same parameters, most electrodes had highly correlated estimated and actual gamma responses, especially for SF-Ori and Con-Ori protocols (Fig. 3B, grey markers). The values were lower for Size-Ori protocol of M2, for which the data points (Fig. 3A, second row) did not smoothly increase with size. This was true of most microelectrodes (Fig. 3B, grey markers), extending from their lower Size-Ori separability (Fig. 2). Interestingly, the same model also predicted ECoG responses reasonably well (Fig. 3B, red markers). The performance using individual electrode parameters (Supplementary Figure 3A) was noticeably better for most ECoGs, but not microelectrodes. The medians were biased towards the microelectrodes because of their higher number compared to ECoGs. Although ECoG and microelectrode tuning preferences tend to be similar^28^, differences in RF size and location may cause some variation in the exact shape of the function. Despite this, the median parameters could represent feature dependencies of ECoG gamma reasonably well. The median parameters obtained by using all trials (not 2 folds) are shown in Table 2.

**Table 2:** Median values of parameters across electrodes obtained by using all trials.

| Parameter | M1 | M2 |
| --- | --- | --- |
| SF center | 5.0196 | 3.0521 |
| SF spread | 0.9413 | 1.0300 |
| ORI center | 109.3960 | 23.9108 |
| ORI spread | 0.8362 | 0.1552 |
| SIZE slope | 0.3674 | 6.3396 |
| SIZE midpoint | 1.2904 | 0.3190 |
| CON slope | 1.4808 | 1.1180 |
| CON midpoint | 22.8935 | 32.1019 |
| HUE center R | 5.9789 | 6.2326 |
| HUE spread R | 1.9068 | 4.2599 |
| HUE center C | 3.0042 | 3.7618 |
| HUE spread C | 2.5095 | 1.8884 |
| HUE gain $a_c/a_r$ | 0.4335 | 0.4030 |
| SIZE slope | 1.6175 | 0.9333 |
| SIZE midpoint | 1.2120 | 0.2257 |
| SAT slope | 3.2922 | 4.2867 |
| SAT midpoint | 0.7477 | 0.6463 |
| VAL slope | 0.0807 | 0.00027 |
| VAL intercept | 0.3472 | 0.4753 |

#### Hue responses

Apart from the stimuli used for separability analysis, we also recorded protocols in which only hue (Hue protocol) or size was varied (Hue-Size protocol). In the Hue protocol, 36 full-screen hues were displayed at maximum saturation and value. Hue-Size protocol displayed red hue at 7 sizes. Hue-Sat and Hue-Val protocols (also used for separability analysis) used 6 hues at 5 levels of saturation and value respectively (see Methods). We first modelled the gamma dependence on individual features as 1-dimensional tuning functions and then used the separability result to express the gamma response as a product of these functions.

The hue response was modelled as the sum of two von Mises functions, each with a center and a spread parameter (Equation 11). The size and saturation were considered as scaling parameters which are maximum at full screen and full saturation. Their tunings were modelled as sigmoidal functions, each with a slope and a midpoint parameter (Equations 12 and 13). The value dependence was modelled as a linear function (Equation 14). We previously found that changing value above 25% caused no change in gamma response^12^ and since we did not sample the 0-25% interval, we skipped the first point (0%) for this fit. The slope of value dependence was close to 0 (the curve was almost flat, “Hue-Val” plot, Fig. 4A), so value had little effect on the gamma response. Reasons for this poor correlation between value and gamma responses are discussed later. As for achromatic case, for all protocols, trials were separated into two folds, parameters were obtained using one fold, and gamma responses from the other fold were correlated against the predicted gamma (see Methods). Parameters obtained for different electrodes are shown in Supplementary Figure 2. Similar to the achromatic case, these values clustered together, so we chose their medians to represent all electrodes in Equation 15A, resulting in the same modelled response for all electrodes except differences in gain and offset. The gain and offset were individually estimated by fitting the function (Equation 15B) with data from the other fold to obtain magnitude matched estimates (Fig. 4A). The linear correlation values between modelled response and data are shown in Fig. 4B. Supplementary Figure 3B shows correlations obtained using individual electrode parameters.

Once the model parameters were fixed for each monkey (Table 2), the gamma response to any achromatic grating specified by four parameters – SF, Ori, Con and Size – was completely determined by equation 10A. Similarly, response to any chromatic patch specified by four parameters – Hue, Sat, Val and size – was determined by equation 15A. Different electrodes could vary only in gains and offsets (Equations 10B and 15B) but that does not change the correlation between actual and predicted gamma responses.

### Prediction of Image Responses

In the natural images protocols, 16 natural images from one of 5 categories were displayed on the full screen (see Methods and ref^4^). One or two categories were displayed in a single protocol. We also used grayscale versions of these images. To predict gamma responses to grayscale images, we checked whether the image patch around the RF of an electrode could be approximated by an achromatic Gabor. For colored images, we tested whether image patches could be represented by a hue patch with relatively uniform HSV values over some radius centered around the RF.

We analyzed grayscale images for grating-like features by approximating them to Gabors having SF, orientation, size and contrast features. We masked the image by a spatial Gaussian of increasing sigma (from 0.3°) and performed a 2-dimensional Fourier transform to identify the dominant SF and orientation within each size before applying some selection criteria (see Methods).However, since natural images have a slow changing feature profile (higher power at low SFs), only a small fraction (1±1.3 out of 16, averaged over electrode- categories) could be approximated with Gabors. We therefore could perform this analysis only for hue patches of colored stimuli, as described below.

For the colored images, we centered discs of iteratively increasing radius (starting with 0.3°) on the RF center, and calculated the means and standard deviations of hue, saturation and value of pixels contained within the enclosed rings. We chose a radius (R) for which these quantities varied within thresholds (see Methods). Patches that did not qualify at the smallest size were assigned a radius of zero. Using pixels contained in R, we calculated their mean hue (H, Equation 16), mean saturation (S, Equation 17) and average value (V) to represent the uniform hue patches. Most images were approximated by at least a small patch. We also tested whether the Value layer could be approximated with a Gabor, as described before.

Images from the Flora category, their magnified sections and approximations for a typical electrode are shown in Fig. 5A. Image 4 was classified as a Gabor based on the Value layer but since it was not achromatic, gamma response could not be predicted from equation 10A. Non-zero radii were obtained for 10 of the remaining images. Using the radius and average hue, saturation and value with Equation 15A gave an estimated gamma response for these 10 images (filled circles in Fig 5C, Selected Set). For images with a patch size of 0 (Images 1, 3, 6-8 here), the model prediction was trivially 0 (Fig. 5C, open circles). The frequency domain response (Fig. 5B) shows gamma peaks for some images. Linear correlation between the actual and estimated gamma response was calculated using all 16 images (rFull=0.58; Fig. 5C) as well as the selected set of 10 images (rSel=0.58). Note that the gain and offset calculation (Equation 15B) would simply change the plot (Fig. 5C) to make the points match in magnitude, but not change the correlations. Also, splitting the data for training and testing is not needed since the model parameters have already been estimated solely using the parametric data.

**Figure 5:**
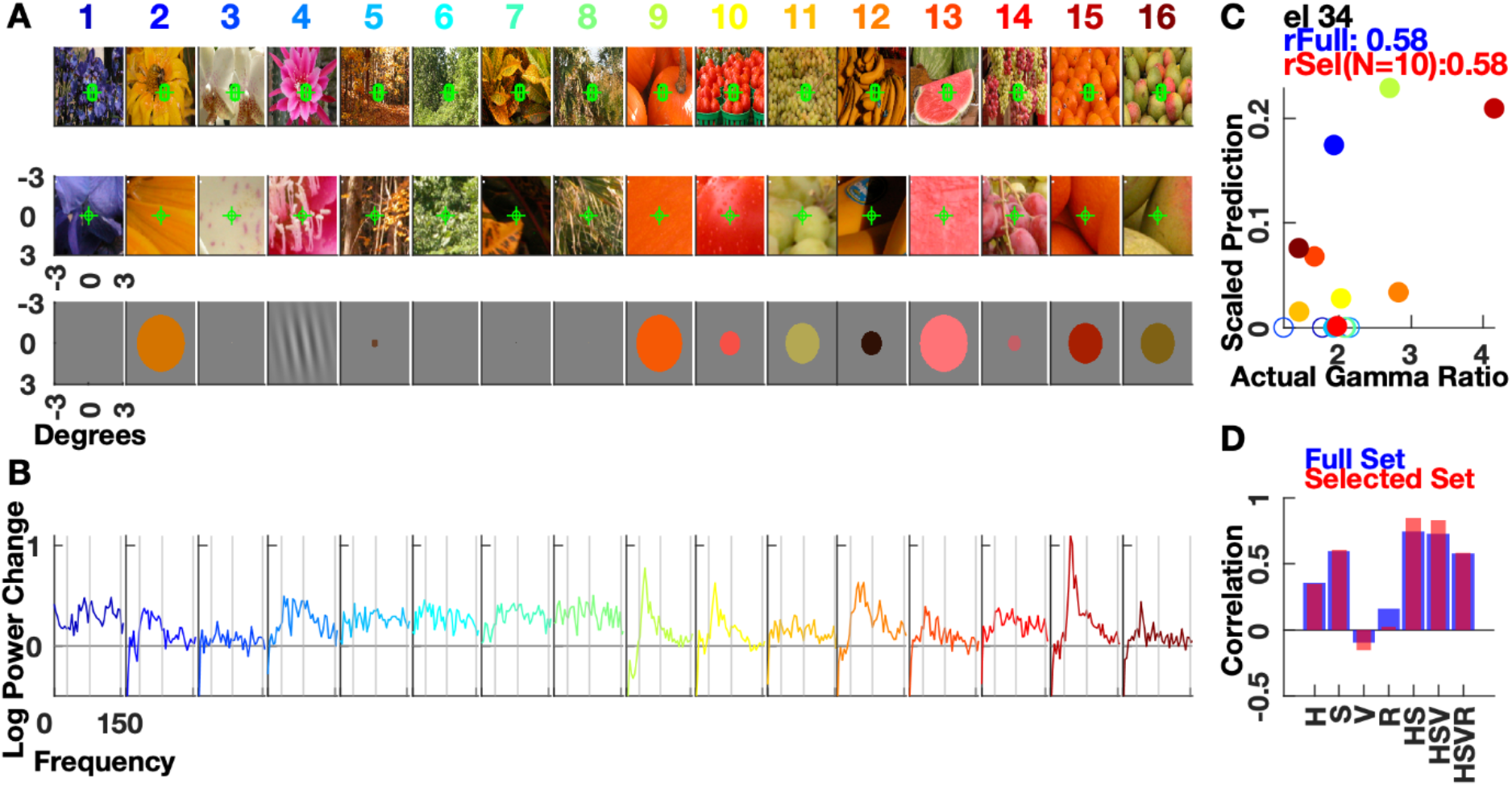
Response predictions for a typical electrode (M1 e34). (A) 16 stimulus images from Flora image set (top row), 3×3 degree patches around the receptive field centre (middle row) and approximations of patches (bottom row). (B) The log change in power in 250-500 ms vs frequency, as obtained by Multi-taper analysis. Vertical lines denote the 30, 80 and 150 Hz frequency marks. (C) Predicted gamma responses versus actual responses i.e. change in power in 30-80 Hz. The filled datapoints correspond to images that are selected as patches in third row of (A). rFull is the correlation for all 16 images, rSel uses only the selected patches. (D) Correlations for full and selected set when chosen patch features (on x-axis: H(ue), S(aturation), V(alue) and R(adius)) were used with the model. The other features were given maximum values, or 0 degrees in case of Hue. The last bar (HSVR) corresponds to data shown in (C).

To check how individual features affect the response estimate, we input only selected features to the model, keeping others constant. For instance, giving only the correct saturation (S), and keeping hue at red (0°), value at maximum (1), and a large size (10°), the correlation across 16 images was 0.59. The correlations for different parameter combinations are shown in Fig. 5D, for both the full and the selected set. The last bar corresponds to Fig. 5C.

### Population Results

We analyzed image patches at all electrode RFs for uniform HSV features. These were used to obtain the response estimate, and the correlation with actual gamma responses was calculated for each electrode and category. The first row in Fig. 6 is the average correlation using different feature combinations in the model (like Fig. 5D). The ‘Full set’ used all 16 images per category (M1:82×5, M2:21×5 (electrodes x categories)). We then dropped patches with size of 0 or a Gabor match to get correlation for the “selected set”. Averaged across electrode-categories, 9.20±2.8SD (of 16) patches qualified as chromatic patches of non-zero size, and 1.13±1.3SD had a Gabor approximation. To eliminate trivial correlations arising due to fewer data points, we only selected electrode-category combinations for which ≥8 (of 16) images qualified as patches (Selected set in Fig. 6).

**Figure 6:**
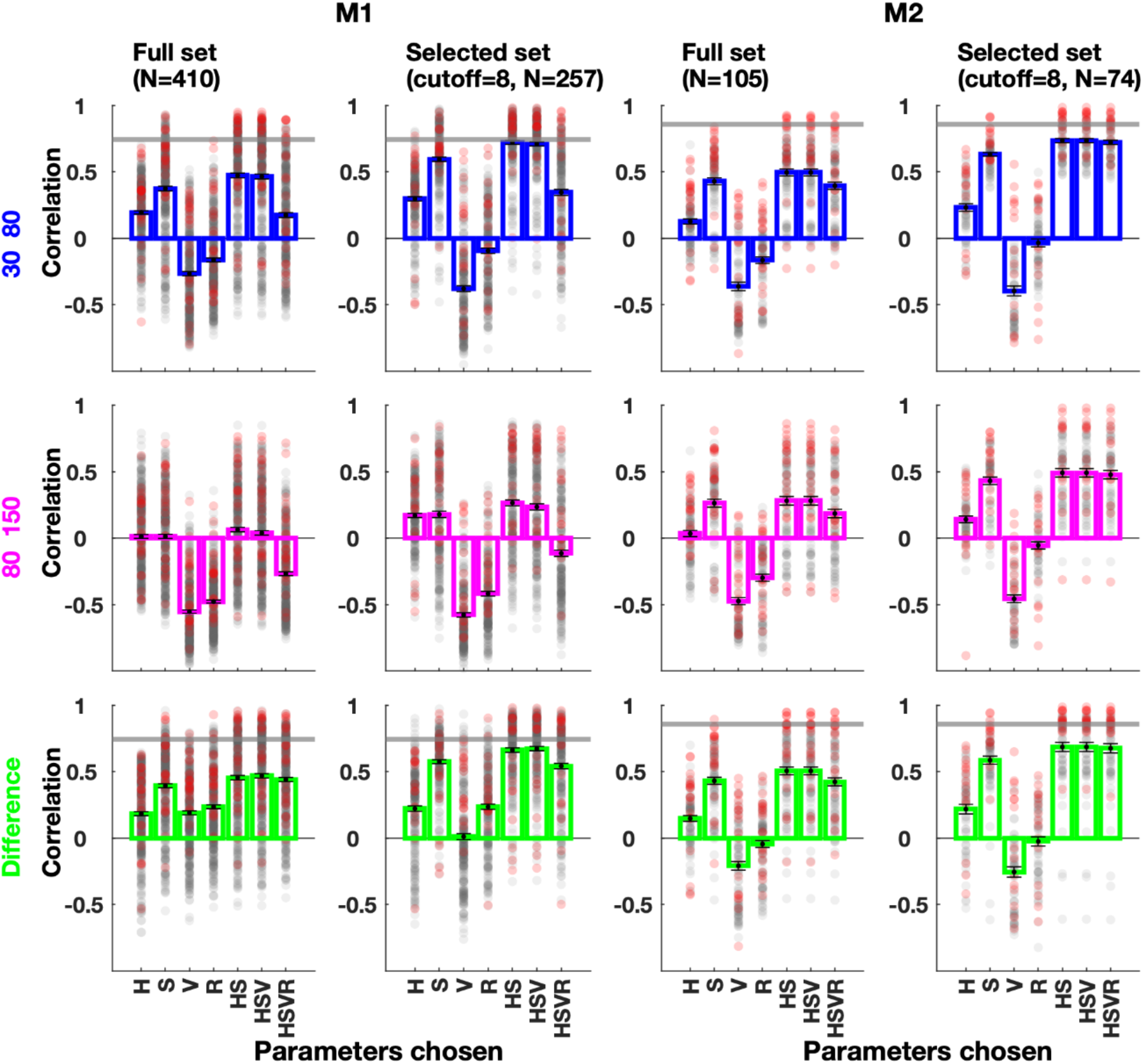
Population Predicted Responses. The correlations between actual response and predicted response modelled with one or more image features. Actual response was chosen as change in power in gamma (30-80, blue) or high gamma (80-150 Hz, magenta) range. The third row (green) is correlation of predicted response with difference between the actual gamma and high gamma response. All sessions (5 image sets for 2 monkeys (82+21 electrodes)) are averaged. In the second and fourth columns, only those sessions are chosen where at least 8 (out of 16) images are selected as patches. Markers show performance of individual electrodes. The horizontal grey lines are the average correlation (across protocols) achieved by the hue model when applied on parametric patches as shown in Fig. 4.

Interestingly, using only value (V) or size (R) resulted in negative correlations. While HS features could predict responses fairly well, addition of R caused a reverse effect, especially for M1. This is because we calculated gamma response as change in power from baseline, but that allows for higher responses even with a wideband pedestal-like increase without an actual gamma peak (Images 5 and 14 versus 9, 10, 15 in Fig. 5B). The reason for this discrepancy is straightforward: image patches tended to be represented as small hue patches when there were salient changes around the RF (e.g. edges), which tended to elicit strong firing. Firing rates have been associated with a broadband spectral response prominent in the high-gamma range^11^. Therefore, small radii as well as dark patches tended to produce higher broadband responses, leading to a negative correlation between V and R parameters and the gamma response.

One solution is to separate the broadband pedestal and the oscillatory response, using recent techniques like FOOOF^31^. However, since the gamma oscillatory response was ambiguous for many images, such separation could depend on the algorithm inputs. We instead followed a simpler strategy: we approximated the wideband response by calculating the change in power in the high-gamma band (80-150 Hz). As expected, the correlations between estimates and high-gamma response were more negative for the V and R only models (magenta bars). To counter the effect of the broadband response, we modified the actual response by taking the difference of the gamma and the high-gamma response. The correlation of this modified response with the estimates (green bars) did not show a steep decrease upon including the size parameter, but did not improve the prediction either.

Overall, the hue and saturation accounted for the most contribution to gamma response, as seen by the HS bars. The correlations for the selected set and HS model were 0.66±0.27SD (M1) and 0.69±0.3SD (M2), while the correlations obtained on parametric hue patches were 0.74±0.21SD and 0.86±0.08SD (averages of points in Fig. 4B, grey lines in Fig. 6). “Value” did not make a significant contribution as expected since gamma versus value was almost flat (Fig. 4A). Size also did not improve the performance partly because the response saturated (Fig. 4A), and partially due to the confounding effect of the broadband response.

The patch parameters depended on the thresholds on the H, S and V parameters: using stringent thresholds lead to smaller radii in general. Since gamma dependence on value was minimal, we had a more relaxed threshold for the V parameter. This can be observed in Images 15 and 16 in Fig 5A, where the estimated patches seem to be larger than expected because the discontinuities were mainly in the “Value” layer. However, since the size did not improve prediction, we also used a basic model in which all patches had a fixed size, over which average H and S were computed as before, and gamma response was estimated using Equation 15A. This basic model also performed reasonably well. For example, for fixed radii of 0.5, 1 and 2 degrees, the respective average correlations for the selected set (same images and electrode- categories as selected with variable radii for Fig. 6) were 0.67±0.26SD, 0.67±0.26SD and 0.66±0.26SD for M1 and 0.66±0.32SD, 0.68±0.30SD and 0.60±0.32SD for M2, as compared to 0.66±0.27SD and 0.69±0.30SD when H and S were computed over variable radii. This shows that even a simple model using only the average hue and saturation of a patch around the RF can estimate the gamma response with reasonably high accuracy.

## Discussion

We found that gamma responses to two stimulus features are largely separable, for both LFP and ECoG scales and could be modelled as a product of individual feature dependencies. We further developed a feature-based image computable model which could predict the gamma responses to chromatic images with reasonably high correlation.

Joint tuning of V1 neuronal responses has been studied for SF and orientation^21, 22, 26, 27, 32^, and features like disparity and orientation or direction selectivity^33^, with varying conclusions about separability. While membrane potentials^24^ and fMRI responses^25^ have been investigated for separability, LFP responses have not. Our results show that the LFP gamma tuning to low-level features of luminance gratings and chromatic patches is largely separable. The separability is maintained across scales as evidenced by the ECoG responses which record from a larger neuronal population. Quantifying the separability also raises questions about data not explained by separable models. This “inseparable component” could result from biophysical or experimental noise, but could indicate other interactions, e.g. subtle shifts in one preferred feature arising from changes in another (as reported in mice^26^), or nonlinear processes contributing to extracellular potentials. Such nonlinearities can potentially be studied via models explaining gamma oscillations through non-linear interactions between excitatory and inhibitory cortical activity^34, 35^. The separability results are surprising, since gamma response was simply the summed power over a fixed (30-80Hz) band, ignoring the gamma peak frequency shifts that occur with stimulus feature changes such as contrast.

We developed a simple model to estimate responses from stimulus features for luminance gratings and hue patches. The model tuning parameters were consistent across electrodes, allowing us to use their medians. Using electrode-wise parameters might improve the model performance (Supplementary Fig. 3), but we traded accuracy for simplicity, generalizing our findings over a cortical area, across scales (LFP and ECoG) and indeed across hemispheres (hue parameters). For the color model, this simplification was necessary since the model was trained on recordings from one brain hemisphere and used to predict responses recorded later from the other hemisphere.

Image-computable models of gamma predict responses to novel images. Earlier attempts in the grayscale space have used luminance contrast as a primary feature. While some studies have shown gamma depends on grayscale image features^16^, others have further predicted the response based on luminance features like orientation^17^, or by using neural networks^19^. Though the effects of color are now being studied, to our knowledge, this is the first attempt at an image-computable gamma model based on chromatic features. Our approach to obtain patches may seem similar to uniformity^16, 18^ and predictability^19^ used by Uran and colleagues. However, we start analyzing the image at the RF whereas they excluded the RF. We also use the OV metric of Hermes^17^ and colleagues; not to calculate gamma but instead as a selection criterion for grating approximations.

Nevertheless, there are several simplifications and limitations of this model. First, the gamma peak frequency is known to vary with stimuli, e.g. increasing with contrast^28^, which we did not take into account. Second, our model does not account for some key drivers of neural activity, such as the luminance change between background and stimulus. Further, the “value” in the HSV space varies between 0-1 for all hues, but corresponds to different luminance for each hue. For example, the interstimulus gray screen (value=0.5) had a luminance of 60cd/m^2^ in our experiments, while the same value (=0.5) for blue, red and green primaries had luminance of 3.5, 13 and 43.5cd/m^2^ (see Table S1 in Ref^12^ for stimulus CIE^36^ (x,y,Y) coordinates; Y is the luminance). While neural activity and gamma response depend on the luminance change, it is not fully captured by the value parameter in HSV space. A better alternative is to convert colors to the physiologically more relevant DKL space^37^, but is beyond the scope here since we did not have responses for stimuli varying along the DKL space. There is some evidence^38^ that when using parametric DKL colors, human MEG gamma is not as strongly biased towards reddish hues, but whether the same holds for V1 LFP gamma responses remains to be tested.

Further, gamma responses to at least two more classes of parametric stimuli are required to properly use our model. The first are achromatic stimuli with mean luminance different from the pre-stimulus gray. Our parametric achromatic stimuli had the same mean luminance of 60cd/m^2^ as the pre-stimulus gray screen (e.g., a 50% contrast grating would have luminance varying between 30 to 90cd/m^2^ across space). However, a natural image can be represented by a 50% contrast grating having a different mean luminance (e.g., a grating whose luminance varies between 10cd/m^2^ and 30cd/m^2^ also has 50% contrast but mean luminance of 20cd/m^2^). When such a grating follows a gray screen of 60 cd/m^2^, the resulting response will be due to two factors– a mean luminance change from 60 to 20cd/m^2^, and a spatial contrast variation of 50%. But gamma dependence on luminance and particularly on the combination of contrast and mean luminance remains unknown. We could have predicted gamma responses to most achromatic images (since they were mainly grayscale uniform patches) had we known the responses for gray at varying luminance, but we did not record this data.

The second stimulus class involves chromatic luminance gratings (like the magenta and white stripes in image 4 of Fig. 5). These are gratings with a particular hue (say red) whose luminance varies with space (generating red-and-black or red-and-white gratings). How the hue, contrast and other factors like orientation combine to produce a gamma response is also unknown. Though previously considered separate pathways, joint coding of color and orientation by V1 neurons has been reported^39^, and may affect the population (gamma) response also. Note that these are different from the iso-luminant chromatic gratings (red-and-green of equal luminance) used in the DKL space. Future studies where these classes of parametric stimuli are also used will allow us to predict a larger set of natural images.

Despite these limitations, we show that a simple model based on mainly the hue and saturation of the pixels inside/around the RF can predict the gamma responses of color images reasonably well. A better understanding of gamma dependence on other classes of parametric stimuli as well as other image-derived measures (like predictability and compressibility) will shed new light on the biophysical mechanisms underlying gamma generated by natural images and their potential role in coding or cognition.

## Materials and Methods

### Animal preparation and recording

Two adult female monkeys (*Macaca radiata*) (M1 and M2 weighing 3.3 and 4 kg respectively) were used in this study. Surgical details have been presented elsewhere^40^ and are briefly described here. The experiments adhered to the guidelines of the Institutional Animal Ethics Committee (IAEC) of the Indian Institute of Science, Bangalore, and the Committee for the Purpose and Supervision of Experiments on Animals (CPCSEA). Customized hybrid arrays were surgically implanted in the left hemisphere of both animals. The custom array had 81 (9 x 9) microelectrodes (Blackrock Microsystems), and 9 (3 x 3) ECoG electrodes (Ad-Tech Medical Instrument Corporation); both connected to a single Blackrock 96 channel connector with common reference wires. The platinum microelectrodes had an inter-electrode distance of 400 μm with a tip diameter of 3-5 μm, and length of 1 mm. The platinum ECoG electrodes had an inter-electrode distance of 10 mm and diameter of 2.3 mm. During surgery, a craniotomy (∼2.8 mm x 2.2 mm) and a smaller durotomy was performed under general anesthesia. The ECoG strip was slid under the surrounding dura (see Fig 1 of Ref^40^). For M2, a part of the strip with 3 electrodes was cut off to assist sliding. A hole made in the silastic was aligned with the durotomy, where the microarray was then inserted into the cortex using a pneumatic inserter. The location of the array was 10-15 mm rostral to the occipital ridge, and 10-15 mm lateral to the midline. Of the ECoGs, 6 (M1) and 4 (M2) were located on V1, posterior to the lunate sulcus. The reference wires were either tucked into the craniotomy crevice or wound around the titanium strap that held the bone flap back in place. The same monkeys had been previously implanted in the right hemisphere V1 with 96 channel Utah arrays^9^ which were not functional by the time the hybrid arrays were implanted. Previously collected recordings from these arrays in response to chromatic stimuli were used as well; and for clarity, the corresponding data are labelled as M1R and M2R.

Following a post-surgery recovery period of ∼10 days, the monkeys regularly performed the experimental tasks. Signals were recorded using Cerebus Neural Signal Processor from Blackrock. LFP and ECoG signals were obtained by band-pass filtering the raw data between 0.3 Hz (analog Butterworth filter, first order), and 500 Hz (digital Butterworth filter, fourth order), and sampling at 2000 Hz. The stimuli were displayed on a gamma corrected monitor (BenQ XL 2411, LCD, 1280 x 720, refresh rate 100Hz). The head fixed monkey viewed the screen from a distance of ∼50 cm. A trial began with the appearance of a fixation dot (0.05° or 0.10° radius) on the gray screen at the center. The monkey passively kept her gaze on the dot and after 1000 ms, 2 - 4 stimuli successively appeared with gray screen in interstimulus period. The fixation dot remained on during the whole trial. Successfully maintaining fixation within 2° of the central dot for the trial resulted in a juice reward, otherwise the trial was aborted.

### Stimuli

#### Achromatic gratings

We studied four features for achromatic gratings – orientation, spatial frequency, size and contrast. Ideally, to study separability of responses, responses to all possible combinations of these parameters need to be recorded. Unfortunately, the number of combinations is prohibitively large in this case. Therefore, we varied only two features at a time – with orientation being the common feature that was varied with one other feature, leading to three combinations as described below. Details of the stimuli and gamma tuning to these features individually has been reported elsewhere^28^.

#### Spatial frequency and orientation stimuli (SF-Ori)

Full screen static gratings at 100% contrast were displayed, at one of 5 spatial frequency values (0.5, 1, 2, 4 or 8 cycles per degree (cpd)) and one of 8 orientation values (0° to 157.5° in steps of 22.5°). The stimuli were randomly interleaved, with 800 ms on and 700 ms off. The stimuli were repeated on average 34.6 (±1SD) and 47(±1.8SD) times for M1 and M2.

#### Size and orientation stimuli (Size-Ori)

The stimulus was a full contrast grating at a given spatial frequency (4 cpd), one of eight orientations (0° to 157.5° in steps of 22.5°) and one of six sizes (0.3°, 0.6°, 1.2°, 2.4°, 4.8° and 9.6° radius). A few sessions of M1 had a seventh size (0.15°), but for the sake of uniformity, it was not used in the analysis here. Stimuli were randomly interleaved and came on for 800 ms, with interstimulus period of 700ms. The stimulus center corresponded to the RF center of one of several electrodes, and varied across sessions (17 sessions for M1, and 7 for M2). On average across sessions, stimuli were repeated 12 (±5.9SD) and 12.6 (±4.1SD) times for M1 and M2.

#### Contrast and orientation stimuli (Con-Ori)

For M1, gratings were displayed at 4 cpd spatial frequency, at one of four orientations (0°, 45°, 90°, 135°) and one of seven contrast values (0, 3.125, 6.25, 12.5, 25, 50 and 100%). The stimuli also had 8 temporal frequency values (between 0 and 50 cycles per second (cps)) and 2 sizes but only the static (0 cps) and full screen case was considered here. For M2, full screen stimuli at 2 cpd spatial frequency were displayed at one of 8 orientations (between 0° and 157.5° in steps of 22.5°) and one of 6 contrasts (0, 6.25, 12.5, 25, 50 and 100%). The mean number of stimulus repetitions was 9.9 (±0.4SD) and 27.9 (±1.6SD) for the two monkeys.

#### Chromatic stimuli

For color stimuli, we used data previously collected using microelectrode arrays from the same two monkeys (right hemisphere, referred to as M1R and M2R), which has also been described in a previous study^12^. We used one protocol in which only hue was varied, one in which only size was varied, and combination protocols in which two parameters from hue, saturation and value were co-varied.

#### Hue stimuli

36 full screen hues equally spaced between 0° (red) and 350° were displayed at full saturation and value for 800 ms with an interstimulus interval of 700 ms. The number of stimulus repeats was 20.05 (±6.2 SD) for M1R and 30.03 (±0.74SD) for M2R.

#### Hue-Size stimuli

A patch of red hue (0°) was displayed at varying sizes, while covering some electrodes of the microelectrode array. For M1R, the sizes were 0.15°, 0.3°, 0.6°, 1.2°, 2.4°, 4.8° and 9.6° and for M2R, sizes were 0.5°, 0.68°, 0.92°, 1.26°, 1.72°, 2.34° and 3.2°. Cyan hue (180°) was also presented for M1R but was not used in the analysis for uniformity. The stimulus came on for 800 ms with interstimulus period of 700 ms. The mean number of repeats of stimuli was 12.4 (±5.2SD) for M1R and 15.9 (±0.4SD) for M2R.

#### Hue-Saturation (Hue-Sat), Hue-Value (Hue-Val) and Saturation-Value (Sat-Val) protocols

In the Hue-Sat protocol, full screen stimuli were presented at six equi-spaced hues (0° to 300°), with saturation values of 0, 0.25, 0.5, 0.75 and 1, while keeping the value at 1 which corresponds to the maximum. At saturation of 0, the screen was white. The stimulus remained on for 800ms with interstimulus period of 700ms. The average number of trials was 12.4 (±0.7SD) and 14.1 (±1.4SD) for M1R and M2R. In the Hue-Val protocol, the saturation was kept fixed at 1, while value could be at 0, 0.25, 0.5, 0.75 or 1 for six equi-spaced full screen hues. At value=0, the screen was simply black irrespective of hue. The mean number of stimulus repeats was 11.8 (±0.6SD) and 27.2 (±0.6SD) for the two monkeys. A Saturation-Value protocol was recorded only for M1R, in which saturation and value both varied in 5 steps from 0 to 1. The stimulus was a large patch (9.6° radius) of red hue (0°) centered at one of the microelectrodes. Each of the 25 stimulus conditions was repeated 11.92 (±4.6SD) times.

#### Image stimuli

Description of the naturalistic stimuli used have been presented earlier^4^. Briefly, 64 images chosen from the McGill Color Calibrated Image Database^41^ (http://tabby.vision.mcgill.ca/html/browsedownload.html) were grouped in 4 sets of 16 each – Fauna, Flora, Textures and Landscapes. We added a set of 16 human face images. All were cropped and down-sampled to get 1280 x 720 sized images. All images are shown in Figure 4C of Ref^4^. We also made grayscale versions of these images using gimp image editor. During a recording session, either 16 (from 1 image set) or 32 (from 2 image sets) stimuli were displayed full screen in a randomly interleaved fashion. Colored and grayscale images were displayed in separate sessions. In each trial, 2-4 full screen stimuli were displayed for 500ms each with an inter stimulus interval of 500ms, during which the monkey maintained fixation. Average number of trials per stimulus across sessions was 72.25(±15.57 SD) for M1 and 69.9 (±5.66 SD) for M2.

#### Post-Processing of Image stimuli

Post experimental recordings, the image stimuli were displayed once again and the screenshots (1280 x 720 pixels) of the display screen were taken. These images were used to obtain image patches centered at the receptive field centers of electrodes. The colored images were in the HSV format, where 3 layers represent the hue, saturation and luminance (value). For the grayscale image, only the luminance layer is relevant, with the hue and saturation set to 0.

### Electrode selection

Electrodes were chosen using a RF mapping protocol described elsewhere^40^. Small gratings were flashed across the visual field and electrodes with consistent responses and reliable RF estimates across sessions were selected. We chose ECoGs which were posterior to the lunate sulcus and had a minimum response value above 100μV. By doing so, we obtained 77 microelectrodes (and 5 ECoGs) in M1, and 17 (and 4 ECoGs) in M2. We also obtained 65 microelectrodes for M1R and 39 microelectrodes for M2R.

In the Size-Ori experiments, stimuli were displayed at RFs of certain electrodes. In each session we selected the electrodes that were within 0.2° of the stimulus center shown in that protocol. We further selected those that showed average firing rate of at least 1 spike/second (250-750 ms), and a signal to noise ratio above 1.5 (details in ref^28^). This yielded 19 unique microelectrodes (and 5 ECoGs) for M1 and 11 (and 4 ECoGs) for M2. Some electrodes were chosen on multiple sessions, yielding 24 and 18 non-unique microelectrodes for the two monkeys. In the Hue-Size protocols, we selected electrodes which were within 0.3° of the stimulus center - 11 in M1R and 35 in M2R. For the Saturation-Value protocol of M1R, 19 electrodes that were within 0.5° of the stimulus center were chosen, for subsequent analysis. The larger range was because the stimulus was also larger in size than Hue-Size protocol.

### Power calculation

Power spectral density (PSD) was calculated by the Multitaper method with 1 Slepian taper, using Chronux toolbox^42^ (https://chronux.org/). The baseline period (spontaneous activity) was 250 - 0 ms before stimulus onset, and stimulus period was 250 - 500 ms after stimulus onset to avoid onset related transients. This led to a frequency resolution of 4 Hz. Power was calculated separately for each trial, and then trial-averaged to get the PSD for each electrode. For the change in power, the stimulus PSD was normalized by the baseline PSD for each electrode. To get the gamma response, PSD values within the 30 - 80 Hz band were summed and then normalized by the summed baseline power in the same range.

### Separability Analysis

The relation between tuning to two features can be characterized by their separability. We tested the separability of gamma responses tuning to two stimulus parameters at a time (say orientation and SF). We used the trial averaged change in gamma band power from baseline as the response to stimuli which varied in the two chosen parameters, resulting in a 2-dimensional tuning matrix with one stimulus feature (orientation) changing across columns and the other feature across rows. This 2-dimensional tuning matrix was expressed as a multiplicative model in two ways - based on Singular Value Decomposition (SVD) and based on marginals^22, 24^.

SVD is a method that allows to decompose a matrix, M in the following format:

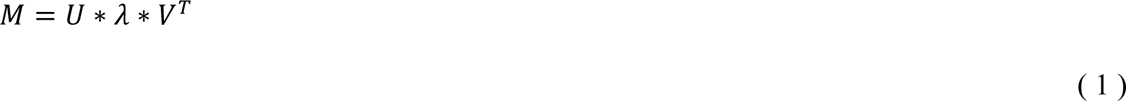

U and V contain orthonormal vectors which are weighted by singular values ordered in the diagonal matrix **λ**. The magnitude of the singular values represent how well M can be approximated as a product of the corresponding singular vectors. The first singular value (λ_1_) corresponds to the largest contribution to the matrix, and so the first singular vectors (U1 and V1) represent the best independent separable factors of M. For a perfectly separable matrix, the other singular values will be zero, which is seldom the case in reality due to measurement noise or biophysical factors. Using the singular values in **λ**, we calculated a separability index^22^ that compares the contribution of the first singular vector to the overall representation:

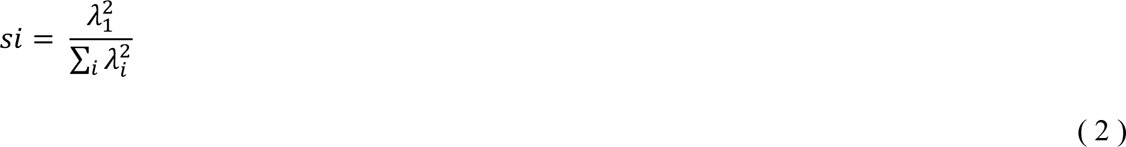

To test the significance of the *si* obtained, we made a 2-dimensional matrix similarly sized as the data with samples randomly chosen from a normal distribution that had the same mean and variance as the joint data matrix, and calculated the *si* of that matrix. We performed 20 such iterations and then tested the hypothesis that the *si* of the data was from the same distribution as that of bootstrapped values, by applying a Student’s t-test (alpha=0.005).

To test how well the first singular vectors represented the data, we used a linear model of the form:

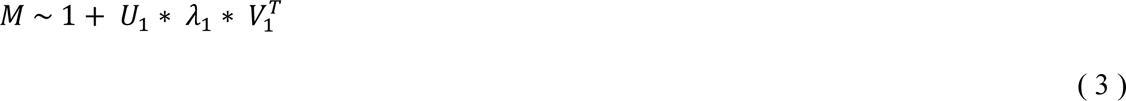

where the first term is an intercept, and the second term is the product of the first singular vectors weighted by the first singular value. We used linear regression to fit this model to data (using the Matlab function *fitlm*) and to obtain coefficients for each of these terms. To evaluate the efficiency of the fit, we calculated the coefficient of determination (R^2^), which represents the percentage of variance of the data that the model can explain and is also equal to the square of Pearson correlation in case of linear regression^43, 44^.

In the second approach, we obtained the marginal responses, F and G from the data matrix by summing across columns and rows respectively. These represent the tuning to one feature (e.g. SF) independent of the other feature (e.g. orientation). For truly independent features, the joint matrix can be reconstructed from the product of marginals in the form:

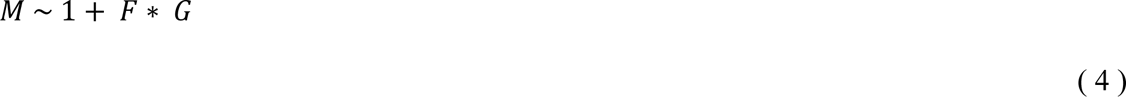

where the first term represents an intercept. We used linear regression to fit the data to this model and obtain the R^2^ metric of goodness of fit. For comparison with the marginal product model, we also tested another linearly separable model (as in ref^24^) based on the addition of the marginals

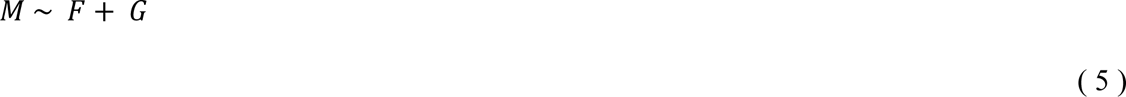

Using linear regression, we obtained the coefficients for all the terms, as well as coefficient of determination (R^2^). There is no intercept here to keep the same degrees of freedom as in equations 3 and 4.

### Modelling of grating tuning functions

We used the responses from the SF-Ori, Size-Ori and Con-Ori protocols to obtain tuning functions of single features and then modelled the joint responses as product of individual tuning functions. In each protocol, trials were divided into 2 non-overlapping folds; trial-average of one fold was used to learn the feature tuning and the other to test it.

The spatial frequency dependence was modelled as a gaussian function over spatial frequency values. Since the SF values doubled as they increased, we applied a log_2_ transformation to linearize them. The orientation tuning was modelled as a von Mises function over the orientation values in radians. Since the grating orientation varied from 0° to 157°, and was symmetrical after 180°, the orientation values were multiplied by 2 to fill the domain of the von Mises function. The 2-dimensional SF-Ori data was used to learn center and spread parameters for the functions:

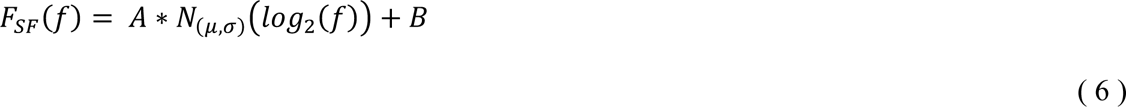

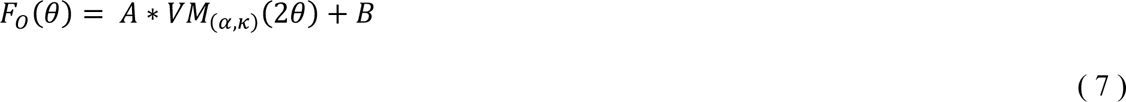

where f and θ represent the spatial frequency (cpd) and the orientation (radians). N_(μ, σ)_ and VM_(α, κ)_ are the Gaussian and von Mises probability density functions normalized by their maximum response, so that the peak value is at 1. By using least squares curve fitting, we obtained the estimates of μ and σ (center and sigma of Gaussian), and α and κ (center and spread of the von Mises curve) as well as gain (A) and offset (B) for each electrode. Each of the two equations has 4 free parameters, which are fitted using 40 data points (5 spatial frequency x 8 orientation values). A and B do not affect the shape of the curve, and merely help to fit the response magnitude.

The effect of size and contrast was modelled as a scaling from the full screen size and the full contrast case. Response was considered to be maximum at the full screen size and at full contrast. The size and contrast values were both log transformed to a linear scale, since they were a geometric sequence. Both were modelled with sigmoidal functions with a slope (*m* for size, *k* for contrast) and a midpoint (*σ_0_* for size, *c_0_* for contrast) determining the shape of the curve. Along with the gain (A) and offset (B), there were 4 free parameters to fit for each of the equations. The fitting was done using data pooled across the orientations to have 48 datapoints (6 sizes x 8 orientations) from Size-Ori data, 28 datapoints (7 contrasts x 4 orientations) from M1 Con-Ori data and 48 data points (6 contrasts x 8 orientations) for M2.

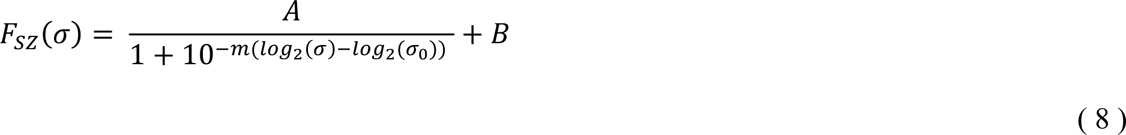

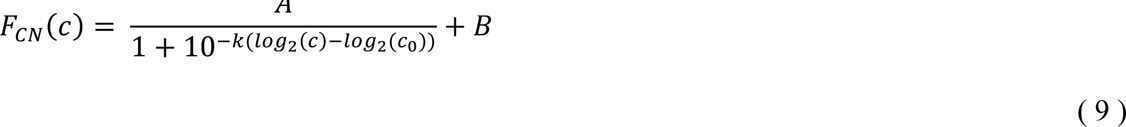

Across the three protocols, we obtained 8 free parameters (leaving out the gains A and offsets B) - μ, σ, α, κ, *m*, *σ_0_, k* and *c_0_* which determined the shape of the tuning for each electrode using one fold of trials. The modelled response to a stimulus was a product of responses to individual features using these parameters.

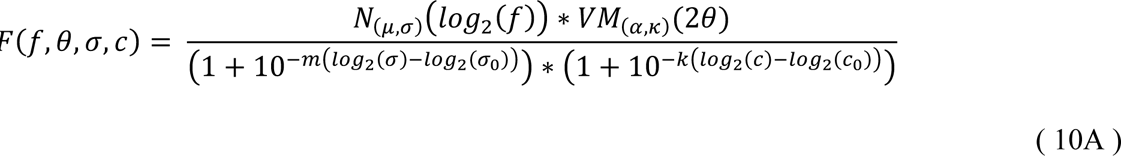

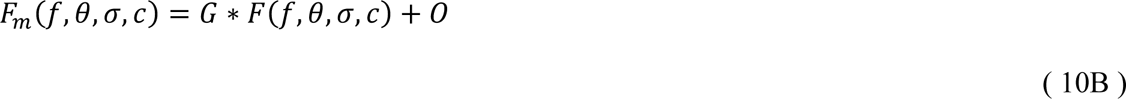

The quantity in Equation 10A depends only on stimulus features and tuning parameters. For the estimates shown in Fig. 3A, the data from the other fold along with the output of equation 10A was used to fit the overall gain G and the intercept O to scale the magnitude of the modelled response (Equation 10B). The linear correlation of this estimate with the actual data was then computed. The process was done in three iterations, so new set of trials would be randomly allotted to each fold. The correlation values were then averaged over the three folds.

### Modelling hue tuning functions

We used responses to the Hue stimuli, Hue-Size stimuli, Hue-Saturation and Hue-Value stimuli, recorded from M1R or M2R to arrive at hue tuning functions. The data was separated into 2 folds, one for obtaining the tuning function parameters and the other for testing. Hue dependence was modelled as a sum of two von Mises functions spread over the circular hue space, with centers initially placed in the region of red (0°) and cyan (180°) hues. The full screen hue response at a chosen hue (H) was modelled as

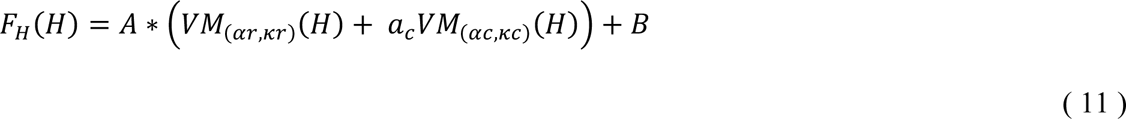

The fitting function had the flexibility to move the centers of the von Mises functions by up to 80° on either side of red, and 100° on either side of cyan center, to cover the whole range of 360°. The gains of the functions were normalized to that at red to be 1 and a_c_. The centers (αr, αc), spreads (κr, κc) and gain (a_c_) of the von Mises functions and the coefficients (A, B) were fit from the Hue data (36 hues), which was supplemented by the maximum saturation and value responses of 6 hues from the Hue-Saturation and Hue-Value stimuli, providing 48 data points to fit 7 free parameters.

The effect of size and saturation was modelled as a scaling on the full screen and full saturation case. The size values were log_2_ transformed to a linear scale. The size was modelled as a sigmoidal dependence as for gratings. It had 4 parameters (slope *m*, midpoint *σ_0_*, gain A, offset B) fit using the size data of red hue (7 data points).

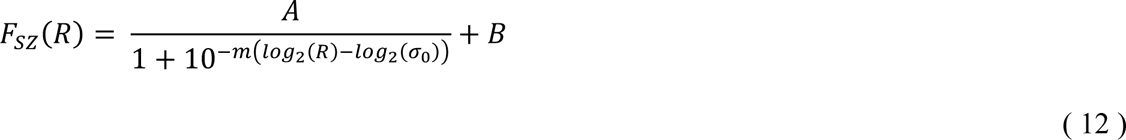

Similarly, saturation tuning was modelled as sigmoidal function with slope *l* and midpoint *s_0_*. These two parameters with the gain and offset were fit using 30 data points (6 hues x 5 saturation) of Hue-Saturation dataset.

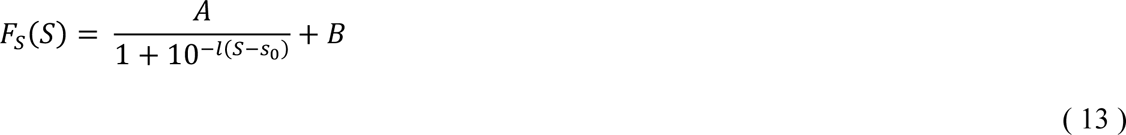

In case of Hue-Value responses, we observed that the gamma remained at around the same value for values 25% and upwards. The model only considered the values above 25% contrast, as we did not have data between 0 and 25%. The values were expressed between 0-1 instead of a percentage. The value dependence was modelled as linear function, with 24 (4 values x 6 hues) data points to fit the gain and intercept:

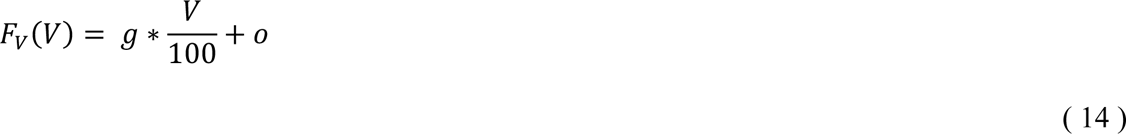

The overall HSV responses were then estimated as a product of individual feature responses:

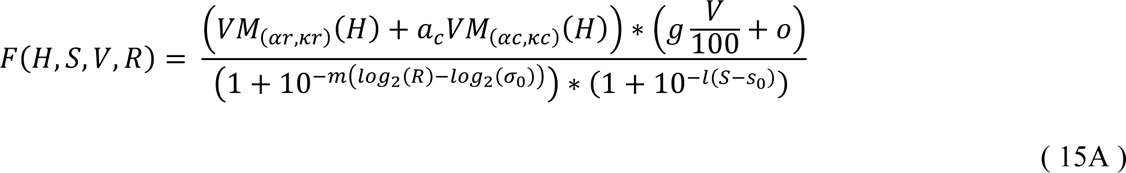

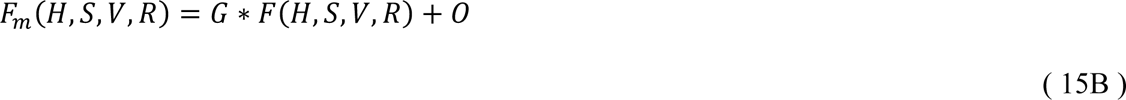

We used the 11 parameters (αr, αc, κr, κc, a_c_, *m*, *σ_0_, l, s_0_, g_v_ and o_v_*) learnt for the individual functions to arrive at the modelled response in Equation 15A. For the results shown in Fig. 4A, these were learnt using one fold of data, and the gain G and intercept O were estimated by fitting this modelled response to the other fold so as to scale the magnitude of the fits (Equation 15B). The fitting of folds was done over three iterations and the correlations between actual and estimated responses were averaged over the three iterations.

The same model (Equation 15A) was used to predict responses to image features. The only difference was that parameters were obtained using the average of all trials, and not from two separate folds as done to estimate the model performance. This did not change the cross validated nature of the analysis, since the image data had no part in calculating the parameters, so all trials of the parametric protocols could safely be used.

### Image patch statistics

We analyzed the image patch around each electrode’s receptive field center for luminance changes similar to gratings, or for unvarying chromatic features. By doing so, most images could be approximated by either a Gabor or a uniform hue patch, which could be defined by simple and clear features.

#### Hue patch approximation

We used all 3 layers of the HSV image centered at the electrodes’ RF. For each electrode- image combination, we calculated the mean (mS_1_, mV_1_) and the standard deviation (sS_1_, sV_1_) of saturation and value in a small patch (0.3° radius) around the RF center. The hues were scaled from 0-360° to 0-2π, and the circular mean (mH_1_) and circular standard deviation (sH_1_) was calculated. For subsequent sizes (increments of 0.3° till 2.1°), we calculated the same quantities (mH_n_, mS_n_, mV_n_, sH_n_, sS_n_, sV_n_), using the pixels in the incremental ring, and excluding the disc of the previous size. We then applied some thresholds on the standard deviations, and the difference of the means from the first size (mH_n_ – mH_1_, mS_n_ - mS_1_, mV_n_ – mV_1_). If any of the standard deviations crossed 0.1, 0.2 and 0.2 for hue, saturation and value respectively, or any of the mean differences crossed 0.05, 0.1 and 0.1, and remained above the threshold for the next two or more radii, we chose that radius as superscribing a uniform HSV patch. These thresholds were chosen because the estimated patch sizes appeared reasonable upon visual inspection, although none of the results shown here critically depend on the choice of the thresholds (as discussed in the Results). If the values dropped to within the threshold range within two steps of crossing, that radius was not chosen, and the next threshold crossing was taken. For the pixels included within the patch of radius R, their average value (V) was calculated to represent the patch. The angular hue (θ) of each pixel was weighted by the corresponding saturation (s) and treated as a vector. The angle and magnitude of their vector sum gave the average hue (H) and saturation (S).

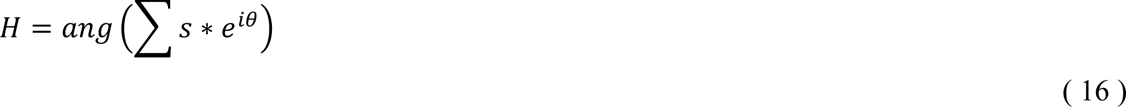

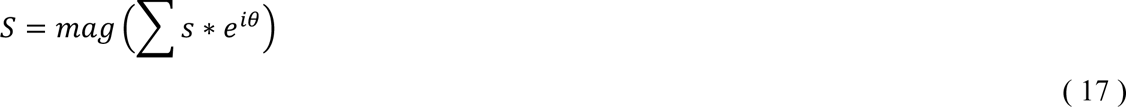

In this way, we got 4 parameters – H, S, V, R – for each image patch. Patches for which the standard deviation threshold was crossed at the first size itself were assigned a size of 0. These values could then be used with Equation 15A to estimate a response.

#### Gabor patch approximation

We used the V layer (of each HSV image) and cropped a square section that was 14° wide around the RF center of each electrode. A Gaussian mask of increasing sigma (starting from 0.3° with increments of 0.3° till 2.1°) was used to highlight the image section within that size. Pixels farther than 3*sigma were set to 0. For the masked image, after removing the DC component, we employed a 2-dimensional Fourier transform to get the magnitude spectra in the spatial frequency and orientation domain. Since naturalistic images have higher power at low frequencies and the spectrum follows a 1/f shape, we chose local maxima in the magnitude spectrum occurring above 0.1 cpd. The local maximum with the highest magnitude was selected and the corresponding spatial frequency and orientation could represent the image at the chosen size. To compare responses across sizes, we normalized the magnitudes with the Euclidean norm of the resulting Gabor at each size. We also obtained the orientation variance (OV^17^) at the given size and SF. OV is the variance in the response magnitudes to different orientated filters at the same SF. We chose the normalized spectral responses for all orientations at the selected SF and calculated their variance. An image patch with a well-defined oriented feature has high OV. Across sizes, we chose the one with the highest normalized magnitude; and in case of a tie, the one with the higher OV. In this way, we got the spatial frequency, orientation and size of the Gabor to match the image. We also estimated the phase and the Michelson contrast of the image in the matched size. We finally selected patches which had SFs between 0.4 cpd and 8 cpd, and an OV above 20. If SF was low, it could be dealt with as a small patch instead of a low frequency grating. If the OV was low, it implied that the chosen orientation was not dominant in the patch, and other orientations could be present, preventing the patch from being well approximated by a Gabor. For the selected images, we obtained the features of spatial frequency (f), orientation (θ), sigma (σ), phase (p) and contrast (c) to make the matching Gabor. As discussed in the Results section, very few image patches qualified as Gabors. This was also observed upon visual inspection of the image patches – only a small minority of images had regularly alternating luminance values across space which would yield a large peak in the 2-D spatial Fourier transform (for example, image 4 in Figure 5A); most images instead were better represented as uniform hue patches of different sizes.

---

**Supplementary Figure 1:**
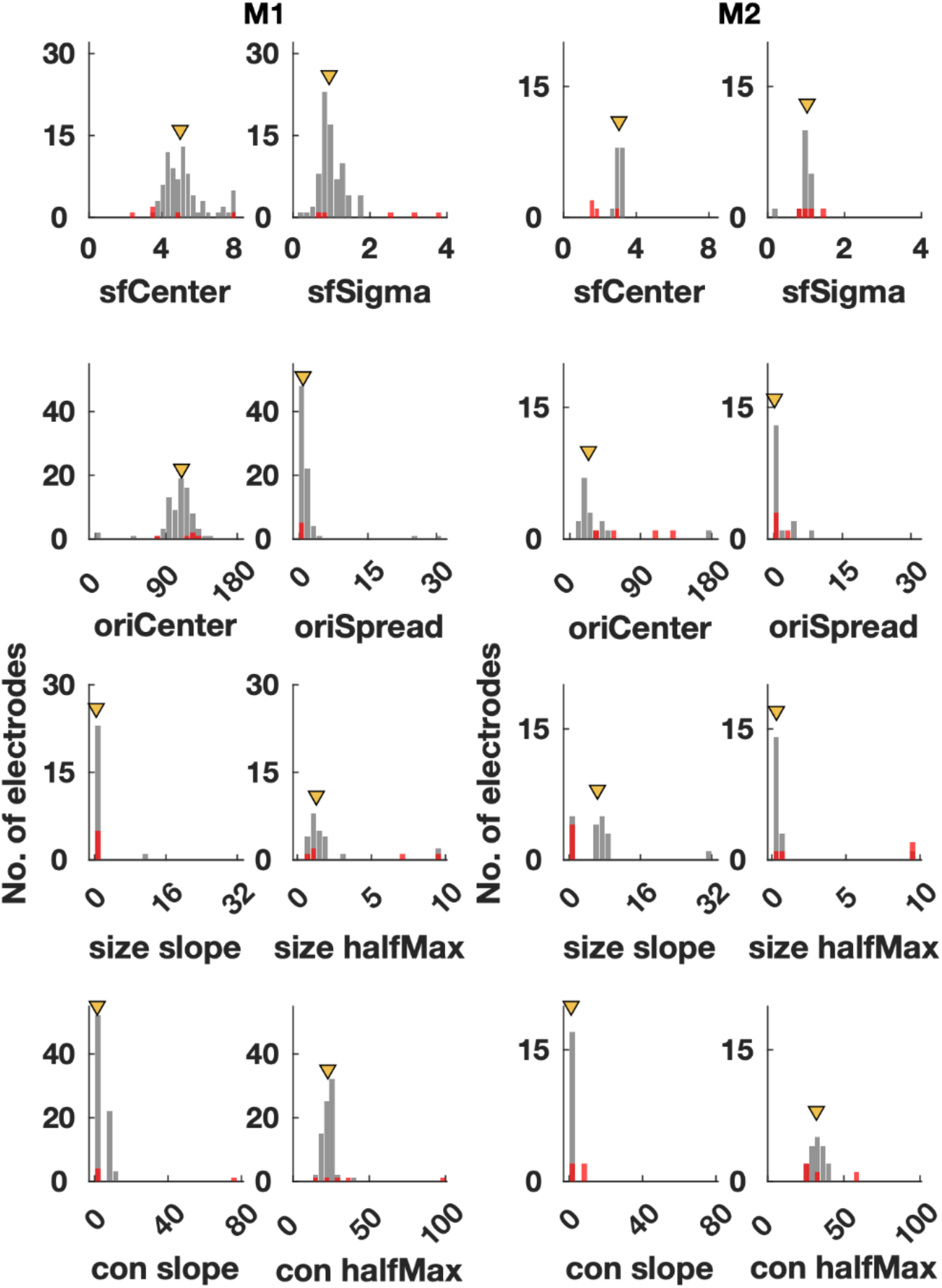
Distribution of grating parameters. Distribution of 8 parameters for all electrodes defining the shape of tuning functions to independent features of spatial frequency, orientation, size and contrast, shown for both monkeys. Note that while trials were divided into two halves for testing the model performance, here these parameters were obtained using all trials (averaging the parameters obtained from two halves also yielded similar results). The grey bars correspond to microelectrodes, and red ones to ECoGs. Medians across all electrodes are shown as triangular markers.

**Supplementary Figure 2:**
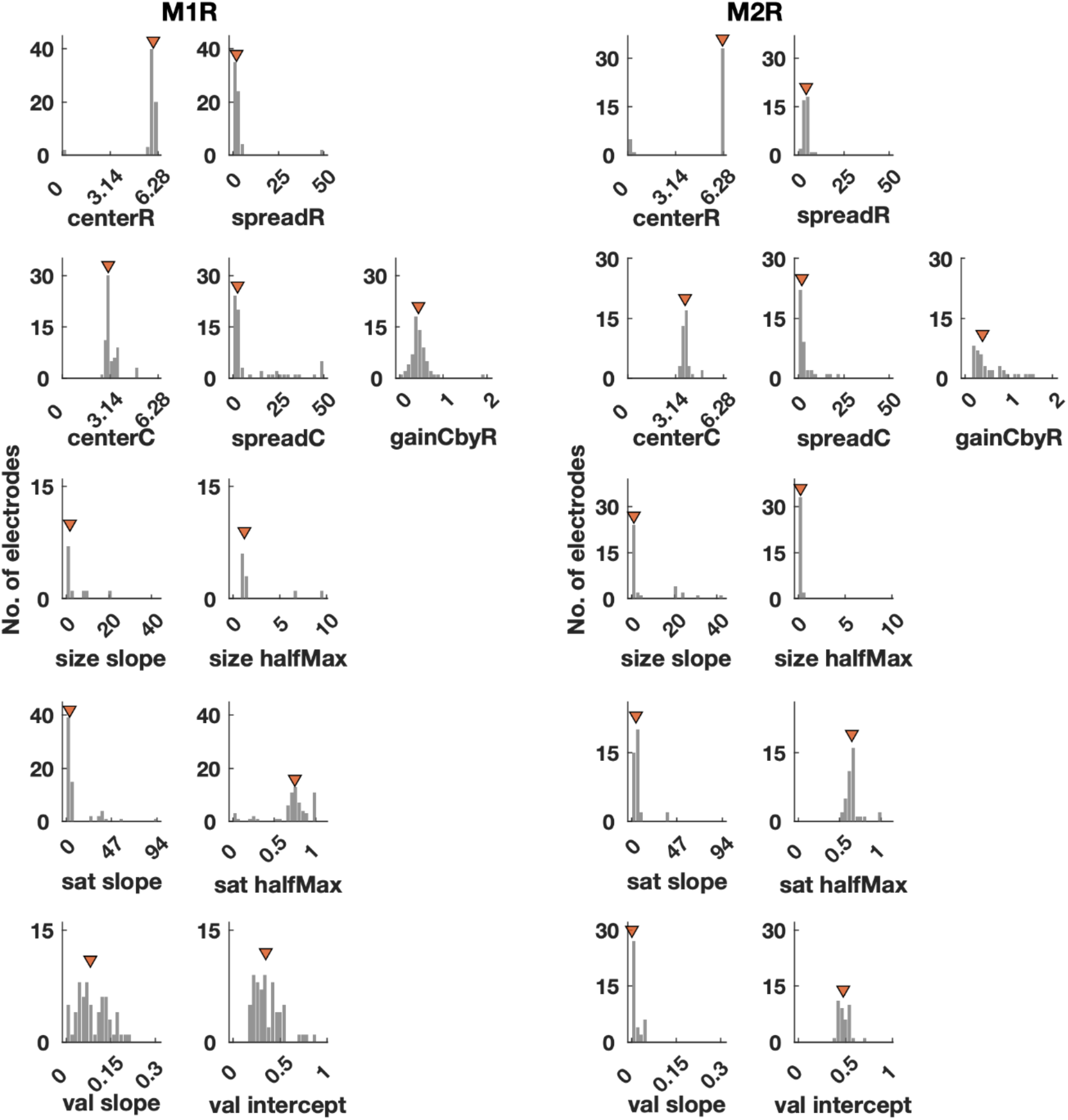
Distribution of hue parameters. Distribution of 11 parameters for all electrodes defining the shape of tuning functions to independent features of hue, size, saturation and value, shown for both monkeys. All trials were used to calculate these parameters. Medians across all microelectrodes are shown as triangular markers.

**Supplementary Figure 3:**
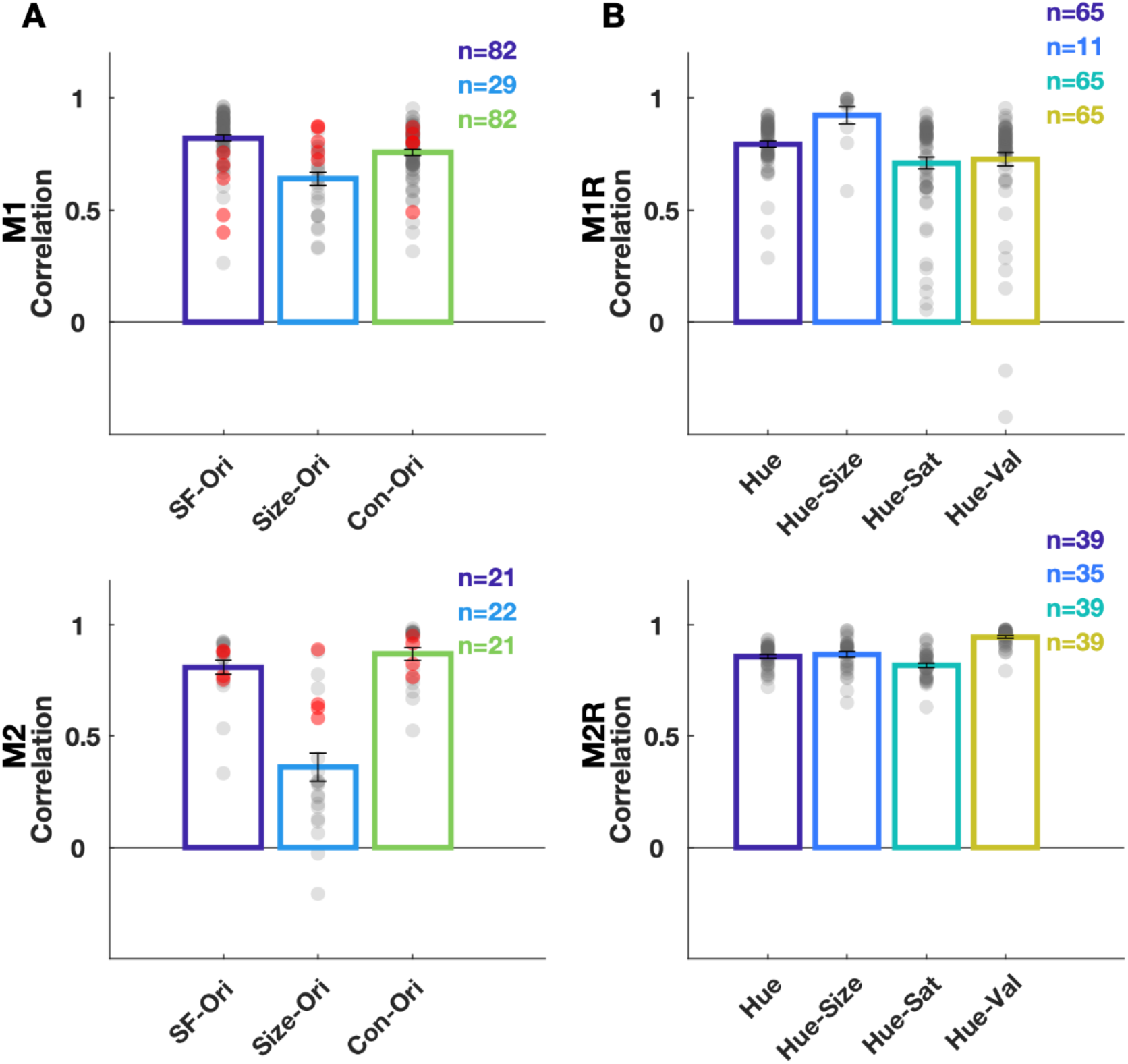
Performance with individual parameters. (A) The correlation between estimated and actual gamma responses when individual tuning parameters of electrodes were used instead of medians, for three grating protocols. Red dots correspond to ECoGs and grey to individual microelectrodes. The correlation values of each electrode are averaged over 3 iterations of cross-validation (B) Correlations between estimates and actual responses for hue protocols, when individual electrode tuning parameters were used. Grey dots represent individual microelectrodes. Error bars correspond to SEM.

## Notes

**Conflict of Interest:** The authors declare no competing financial interests.

**FUNDING DISCLOSURE:** This work was supported by DBT/Wellcome India Alliance (IA/S/18/2/504003; senior fellowship to SR) and a grant from Pratiksha Trust.

### Competing Interest Statement

The authors have declared no competing interest.

